# Runaway evolution from male-male competition

**DOI:** 10.1101/2021.05.17.444494

**Authors:** Allen J. Moore, Joel W. McGlothlin, Jason B. Wolf

**Affiliations:** Department of Entomology, University of Georgia, Athens, Georgia, United States; Department of Biological Sciences, Virginia Tech, Blacksburg, Virginia, United States; Milner Centre for Evolution and Department of Biology and Biochemistry, University of Bath, Bath, United Kingdom

**Keywords:** aggression, honest signals, indirect genetic effects, male-male competition, quantitative genetics, runaway evolution, sexual selection, social evolution, weapons

## Abstract

Wondrously elaborate weapons and displays that appear to be counter to ecological optima are widespread features of male contests for mates across the animal kingdom. To understand how such diverse traits evolve, here we develop a quantitative genetic model of sexual selection for a male signaling trait that mediates aggression in male-male contests and show that an honest indicator of aggression can generate selection on itself by altering the social environment. This can cause selection to accelerate as the trait is elaborated, leading to runaway evolution. Thus, an evolving source of selection provided by the social environment is the fundamental unifying feature of runaway sexual selection driven by either male-male competition or female mate choice. However, a key difference is that runaway driven by male-male competition requires signal honesty. Our model identifies simple conditions that provide clear, testable predictions for empirical studies using standard quantitative genetic methods.

## INTRODUCTION

Darwin (1859, 1871) chronicled the amazing diversity of traits associated with success in mating, proposing the theory of sexual selection to explain the diverse exaggerated, spectacular, and bizarre structures and behaviors found in males of many species. Darwin suggested that such traits evolve either because they enhance success in contests between males for access to females or because they are preferred by females when choosing mates. As Darwin (1871) wrote, “It is certain that amongst almost all animals there is a struggle between the males for the possession of the female. This fact is so notorious that it would be superfluous to give examples.” In contrast, the ability of females to influence evolution through choice of partners was almost immediately questioned and continued to be controversial for decades after Darwin (Wallace 1889; Huxley 1938). However, theoretical models of evolution via female choice (Lande 1981; Kirkpatrick 1982; Mead & Arnold 2004) and empirical research documenting female preference in nature (Andersson 1982, 1994; Rosenthal 2017) eventually led to mate choice becoming the dominant paradigm in studies of sexual selection. The development of formal mathematical models showing that male traits and female preferences coevolve in self-reinforcing fashion, an idea first proposed by Fisher (1915, 1930), was particularly crucial to the acceptance of mate choice as an important evolutionary mechanism. The key component of the Fisher process is that female preference and a preferred male trait become genetically correlated as a result of assortative mating that generates linkage disequilibrium between the preference and male trait. This can cause sexually selected male traits to evolve at ever-increasing speed, a pattern that has been referred to as an evolutionary “runaway” (Fisher 1930; Lande 1981; Bailey & Moore 2012).

Despite the current bias towards studies focused on mate choice, Darwin was not wrong about male-male competition. Members of entire taxa are characterized by highly modified sexually dimorphic structures that function only in male contests (e.g., Dermaptera, Briceño & Eberhard 1995). Weapons can evolve to be massive and create real functional constraints for the males that bear them, and such bizarrely elaborate and diverse structures associated with duels are indeed found across the animal kingdom (Emlen 2008, 2014; McCullough *et al.* 2016; O’Brien *et al.* 2018). In fact, male-male competition remains a more common source of selection shaping male traits that influence mating success, and traits expressed in male-male interactions can be as elaborate as those that are the target of female preferences (Darwin 1871; Huxley 1938; Andersson 1994; Moore & Moore 2006; Emlen 2008, 2014; McCullough *et al.* 2016; O’Brien *et al.* 2018). However, we still lack robust genetic models that generate testable predictions for the evolution of sexually selected traits via male-male competition. Notably, the potential for male-male competition to result in a runaway process that drives extreme trait elaboration remains unresolved.

Many elaborate male traits used in male-male contests, such as showy plumage (Hagelin 2002), color (Seehausen & Schluter 2004), pheromones (Moore *et al.* 1997b), and structures such as antlers (Wilkinson & Dodson 1997), horns (Emlen *et al.* 2005), forceps (Briceño & Eberhard 1995), and claws (Sneddon *et al.* 1997) function as signals that may provide information about some underlying qualities of the individuals, such as the willingness or ability to fight (Parker 1974; Maynard Smith & Harper 1988; Maynard Smith & Harper 2003; Emlen 2008, 2014). For example, there is often a positive association between signals or weapons and other traits such as body size (McCullough *et al.* 2016; O’Brien *et al.* 2018), making the signal or weapon an honest indicator of potential threat to an opponent. As such, males are expected to adaptively modulate their behavior in response to these signaling traits, escalating contests they are more likely to win and withdrawing from ones they are more likely to lose. Because the effect of signaling traits inherently depends on social context, such traits serve as both targets and sources of selection, potentially leading to self-reinforcing and accelerating selection as occurs in the runaway process driven by female preference (Lande 1981; Bailey & Kölliker 2019). However, despite insights from game theory models (Parker 1974; Maynard Smith & Brown 1986; Maynard Smith & Harper 1988; Maynard Smith & Harper 2003; Rutte *et al.* 2006), how this fundamental feature of extreme elaboration – an evolving source of selection – may arise within male-male contests is unclear.

Here, we utilize a framework that explicitly incorporates socially contingent trait expression and fitness (Moore *et al.* 1997a; Wolf *et al.* 1999; McGlothlin *et al.* 2010) to model trait evolution arising from male-male competition. We show that when honest signals are used to modulate the behavior of competitors, male-male competition leads to evolutionary elaboration of male traits. We identify the necessary and sufficient conditions for trait elaboration to become a runaway process and outline predictions that can be empirically tested to evaluate this scenario in natural systems. We show that sexual selection by male-male competition can have features that are analogous to those of runaway sexual selection by female choice; just as in female mate choice, the social environment in male-male contests may generate a self-reinforcing source of selection on the traits that mediate the interaction, potentially leading to self-sustaining and escalating selection.

## MODEL

To capture the influence of the social environment in a model of male-male competition, we assume that individuals adjust their behavior in response to the signaling trait values of their social partners, an assumption that is supported empirically and theoretically (Parker 1974; West-Eberhard 1979; Maynard Smith 1982; West-Eberhard 1983, 1984; Moore *et al.* 1997a; West-Eberhard 2003; O’Brien *et al.* 2018; Tinghitella *et al.* 2018; Rico-Guevara & Hurme 2019; Wiens & Tuschhoff 2020). Because the social context (i.e., the social environment) is constructed from traits of conspecifics, this flexible response to social context provides the opportunity for indirect genetic effects (Moore *et al.* 1997a), which allow the social environment itself to evolve (Moore *et al.* 2002; Wolf 2003; Bijma & Wade 2008; McGlothlin *et al.* 2010). Evolutionary changes in the social environment can lead to concerted evolution because the social environment can be a source of selection on the traits that themselves compose the social environment (West-Eberhard 1979; Wolf *et al.* 1999; Bailey & Kölliker 2019; Araya-Ajoy *et al.* 2020; McGlothlin *et al.* 2022). Such “social selection” (West-Eberhard 1979, 1983, 1984; Wolf *et al.* 1999; Bijma & Wade 2008; McGlothlin *et al.* 2010) is expected to arise whenever traits act as both agents and targets of selection.

Our model assumptions are based on common conditions observed in male-male contests (Emlen 2008, 2014; Eberhard *et al.* 2018). Although we model the outcome of pairwise duels between males drawn at random from the population, our model is easily generalized to include multiple interactions between males (Supporting Appendix). First, we assume that males possess a trait (designated by the subscript *S*) that is used as a signal conveying potential threat in social contests. There are many diverse examples include traits such as plumage patches, exaggerated weapons, or vocal or chemical signals. Elaboration of the signal may consist of an increase in size or complexity, although for heuristic simplicity, we discuss the evolution of increased signal size. Second, we assume that males vary in the underlying quality trait that reflects their fighting ability or some other aspect of their phenotype that determines the potential interaction cost they represent to their opponent in a contest. We describe this trait as body size (designated by the subscript *B*) for simplicity (see the discussion of male quality in Eberhard *et al.* 2018). Finally, we assume males respond to the assessment of the signal by modulating their behavioral response of aggression toward their opponent (designated by the subscript *A*) within the contest because the signal provides information on the likelihood that they would win an escalated contest (see below).

We assume that both signal size (*Z*_*S*_) and body size (*Z*_*B*_) are normally distributed metric traits influenced by many loci of small effect. Expression of these traits can be partitioned into heritable additive genetic effects (denoted *a*_*S*_ and *a*_*B*_) and general non-heritable (environmental and nonadditive genetic) effects (denoted *e*_*S*_ and *e*_*B*_). We assume that neither signal nor body size changes as a result of the social interaction. An individual’s total phenotypic value for each trait is then described by a simple sum of the heritable and non-heritable components:

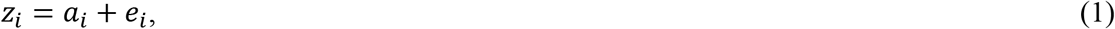

where *a*_*i*_ is normally distributed with mean 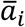 and variance *G*_*ii*_ and *e*_*i*_ is normally distributed with mean 0 and variance *E*_*ii*_ We make the standard quantitative genetic assumption that heritable and non-heritable components are uncorrelated.

We model male-male competition where the larger males will defeat smaller males in a fight. We therefore further assume that the phenotypic value for aggressive behavior (*Z*_*A*_) associated with a given genotype depends on social context, influenced by their rival’s signal size relative to their own, as suggested by West-Eberhard (1979, 1983, 1984). This effect is captured in our model as a term in which aggression scales with the magnitude of the size difference between opponents, which is supported by optimality models and empirical studies (Huxley 1938; Parker 1974; Riechert 1984; Sneddon *et al.* 1997; Maynard Smith & Harper 2003; Emlen 2008, 2014). The phenotypic value of aggression can thus be written:

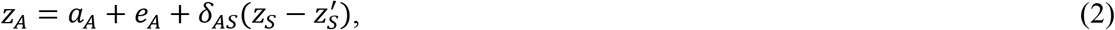

where *a*_*A*_ and *e*_*A*_ represent standard additive genetic and uncorrelated non-heritable components, respectively. Here and elsewhere, terms with primes indicate a value assigned to the focal individual’s opponent so *Z*_*S*_ is the phenotypic value of the signal of the focal individual and 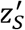 of their opponent. The coefficient *δ*_*AS*_ measures the influence of the difference in signal size on the expression of aggressive behavior. Thus, *δ*_*AS*_ is analogous to the *ψ* term in standard interacting phenotype models (Moore *et al.* 1997a), but differs because it depends upon the value of an interactant’s phenotype relative to the focal individual. Because signal size is heritable, the phenotype value of aggression for the focal individual includes modifications arising from both direct genetic effects of their own genotype (*δ*_*AS*_*a*_*S*_) and indirect genetic effects 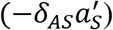, which is defined as the effect of a social interactant’s genotype on the focal individual’s phenotype (Moore *et al.* 1997a). We describe the relationship between this model and the standard model of indirect genetic effects in the Supporting Appendix.

The underlying genetic value of aggression is assumed to be genetically uncorrelated to both that of signal size and body size (i.e., there is no direct pleiotropic relationship between the traits such that genetic covariances *G*_*SA*_ = *G*_*BA*_ = 0). This is a conservative assumption as a positive correlation would result in even faster evolution. However, the signal may be genetically correlated to body size, providing signal honesty, which is quantified by the covariance between signal size and body size (*G*_*SB*_). Because the level of aggression displayed is conditional on the social context, a correlation within the population is generated if males with larger signals and/or larger body size are more aggressive on average (and vice versa). Hence, aggression can be correlated to the signal and body size traits despite the absence of a direct pleiotropic link (or linkage disequilibrium) because the flexible behavioral response creates a relationship between these traits through the social interaction.

### Selection imposed by male-male competition

In social interactions, associations between traits and fitness may cause selection via two pathways: nonsocial selection (quantified by the gradient *β*_*N*_), which arises from effects of a focal individual’s traits on its own fitness, and social selection (quantified by the gradient *β*_*S*_), which arises from the effects of an opponent’s traits on the fitness of a focal individual (Wolf *et al.* 1999). From Wolf *et al.* (1999), when both nonsocial and social selection are present, individual relative fitness can be written:

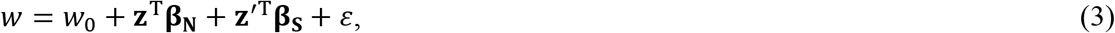

where *w*_0_ is an intercept, **z** and **z**′ are column vectors of focal and opponent traits, **β**_**N**_ and **β**_**S**_ are vectors of nonsocial and social selection gradients, *ε* is an uncorrelated error term, and the superscript T denotes transposition. Expressing relative fitness using Eq. 3 has two distinct advantages. First, selection gradients can be estimated in natural populations using multiple regression (Lande & Arnold 1983; Wolf *et al.* 1999; Formica *et al.* 2011; Fisher & Pruitt 2019), allowing our model to generate testable predictions. Second, selection gradients can be combined with genetic parameters to predict short-term evolutionary response to selection (Lande & Arnold 1983; Bijma & Wade 2008; McGlothlin *et al.* 2010).

To understand how these selection gradients arise from male-male contests with signaling, we can use evolutionary game theory (see Supporting Appendix) to write a mechanistic expression for relative fitness:

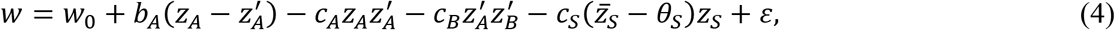

where terms including *b* represent fitness benefits and terms including *c* represent fitness costs. In Eq. 4, the benefit term and the first cost term derive from the hawk-dove model of evolutionary game theory (Appendix; Maynard Smith & Price 1973; Maynard Smith 1982; McGlothlin *et al.* 2022). The coefficient *b*_*A*_ is the fitness benefit of winning a contest, which we assume derives from greater access to females. In a contest, access to females is determined by a focal individual’s aggression relative to its opponent. Multiplying *b*_*A*_ by 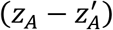 reflects the fact that the probability of winning a contest increases as a male can become increasingly more aggressive than its opponent. This benefit, however, does not come without a cost. The term 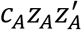 is the fitness cost of aggression associated with escalation of encounters. Logically, an individual pays a cost for acting aggressive that depends on the level of aggression shown by its opponent. In the hawk-dove model, this corresponds to the cost associated with a hawk player facing a hawk opponent, which increases in likelihood as both players act increasingly aggressive. A second fitness cost 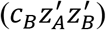 reflects the fact that the fitness impact of aggression by an opponent 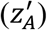 depends on the size of the opponent 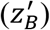 This cost, which we call the threat of the opponent, derives from the fact that larger males impose a greater risk of harm than do smaller males. Finally, we assume that a third cost 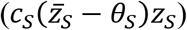 arises from natural selection favoring some optimal trait value (*θ*_*S*_), which therefore will oppose signal elaboration. Following a Gaussian model of selection (Lande 1976, 1979), selection against elaborate signals becomes stronger as the population mean of the signal 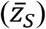 becomes further away from its naturally selected optimum (*θ*_*S*_). Although we do not do so here, this term could be replaced with a multivariate Gaussian term (Lande 1979) to add naturally selected optima for aggression and body size.

Taking partial derivatives of with respect to focal and opponent traits (evaluated at the population mean) allows us to translate the fitness model in Eq. 4 into nonsocial and social selection gradients (McGlothlin *et al.* 2022; Supporting Appendix). The nonsocial gradients are:

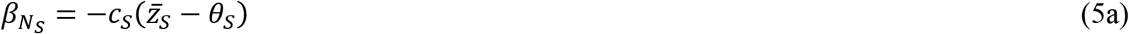

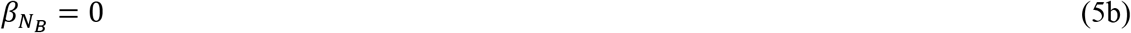

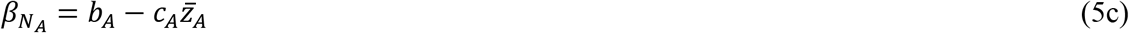

and social selection gradients:

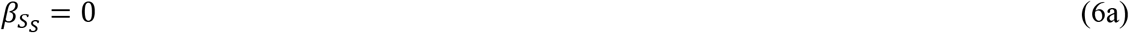

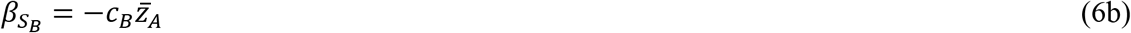

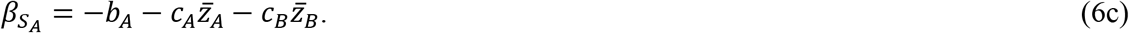

Thus, males with large signals are selected against via nonsocial selection but interacting with such males does not directly impose social selection (Eq. 5a). Body size is not under direct nonsocial selection but imposes a fitness cost via social selection that increases with the population mean of aggression (Eq. 5b). Nonsocial selection favors aggression until the benefits of aggression are outweighed by the costs, while social selection imposed by opponent’s aggression is always negative, representing a net fitness cost (Eq. 5c). This gradient becomes increasingly negative as the population mean aggression and body size increase. These selection gradients suggest that signal size itself experiences no direct sexual selection. If signal size increases, it must do so as an indirect response to selection on a correlated trait.

### Evolutionary response to selection

Selection within a generation is translated into an evolutionary response across generations through the association between the phenotype, upon which selection acts, and the genotype, which contributes to the inheritance of the traits across generations. In quantitative genetics, this genotype-phenotype relationship is most often summarized by the additive genetic variance, which is used to predict evolutionary response to selection across generations (Lande & Arnold 1983; Arnold 1994). However, for traits expressed in social interactions, we must also consider social pathways to fitness, which arise from indirect genetic effects and social selection, when calculating response to selection (Moore *et al.* 1997a; Bijma & Wade 2008; McGlothlin *et al.* 2010). Because the model of phenotypic modification described in Eq. 2 deviates from the standard model of indirect genetic effects, we develop a general equation for response to selection in the Supporting Appendix (Eq. A10). Using this equation, the response to selection for the three traits in our model can in general be written:

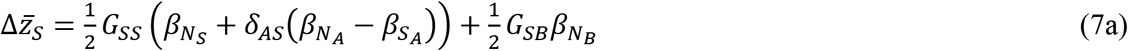

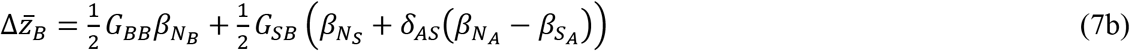

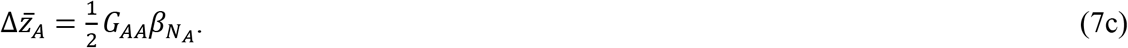

The multiplier ½ in Eq. 7 arises because selection is acting only on males. Eq. 7a shows that modification of aggressive behavior in response to the signaling trait (*δ*_*AS*_) causes both nonsocial and social selection gradients for aggression to contribute to signal evolution. This behavioral modification also contributes to evolution of body size when the signal is honest, which is captured by the covariance between signal size and body size (*G*_*SB*_; Eq. 7b). This is easily shown by setting the modification of aggression based on the signaling trait (*δ*_*AS*_) to 0, which recovers standard quantitative genetic expressions for evolution. In contrast, behavioral modification never contributes to evolution of aggression (Eq. 7c).

By substituting Eqs. 5-6 into Eq. 7, we can predict evolutionary change using our mechanistic fitness model (Eq. 4):

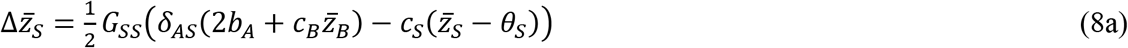

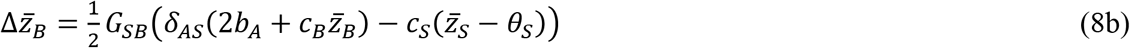

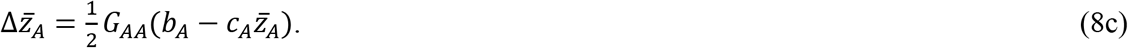

Eq. 8a shows that when fitness is defined as in Eq. 4, evolution of the signaling trait beyond its naturally selected optimum depends crucially on modification of aggression. If males do not change their aggression to the signal (i.e., if *δ*_*AS*_ = 0), the population mean of the signaling trait cannot increase. From Eq. 8b, the evolution of body size depends on both *δ*_*AS*_ and the presence of signal honesty (i.e., *G*_*SB*_ > 0). Eqs. 8a-b also show that evolution of the signaling trait and of male body size is potentially open-ended because the evolutionary response to selection for each trait becomes stronger as the population mean body size increases. In contrast, from Eq. 8c, the evolution of aggression is self-limiting because selection depends on the balance of the benefits and costs of aggression, the latter of which become more intense as mean aggression intensifies. This observation suggests that both signal size and body size may experience runaway evolution if the benefits of aggression and the threat of the opponent are strong enough to outweigh natural selection against elaborate signals, whereas aggression should always tend to quickly evolve to an equilibrium value.

To solve for equilibrium and to explore the conditions for such a runaway, we set Eqs. 8a-c equal to zero and solve for the equilibrium mean of each trait 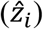:

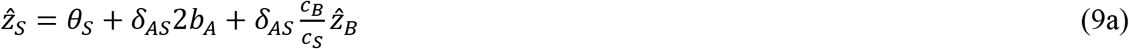

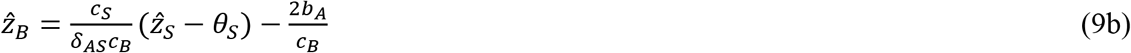

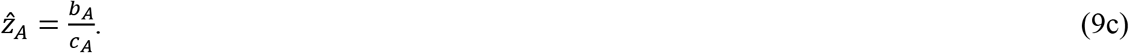

As predicted, aggression will always reach a stable equilibrium whenever there is a cost of aggression (Eq. 9c, Fig. 1). Eqs. 9a-b predicts a line of equilibria for signal size and body size, because their evolutionary change is completely intertwined with the relationship 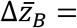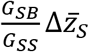 (Fig. 1). The slope of the line of equilibria predicting mean signal size from mean body size, and hence the evolutionary allometry of signal size, is 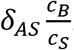. This relationship indicates that when comparing population means through time or across space, positive allometry (i.e., a slope greater than unity) is predicted when the strength of behavioral modification multiplied by the threat of the opponent (*δ*_*AS*_*C*_*B*_) is greater than the strength of natural selection on signal size (*C*_*S*_). In general, when male behavior is more strongly dependent on the signal of their opponent (i.e., when *δ*_*AS*_ is larger), more elaborate signals are expected at equilibrium (Fig. 2).

**FIGURE 1.**
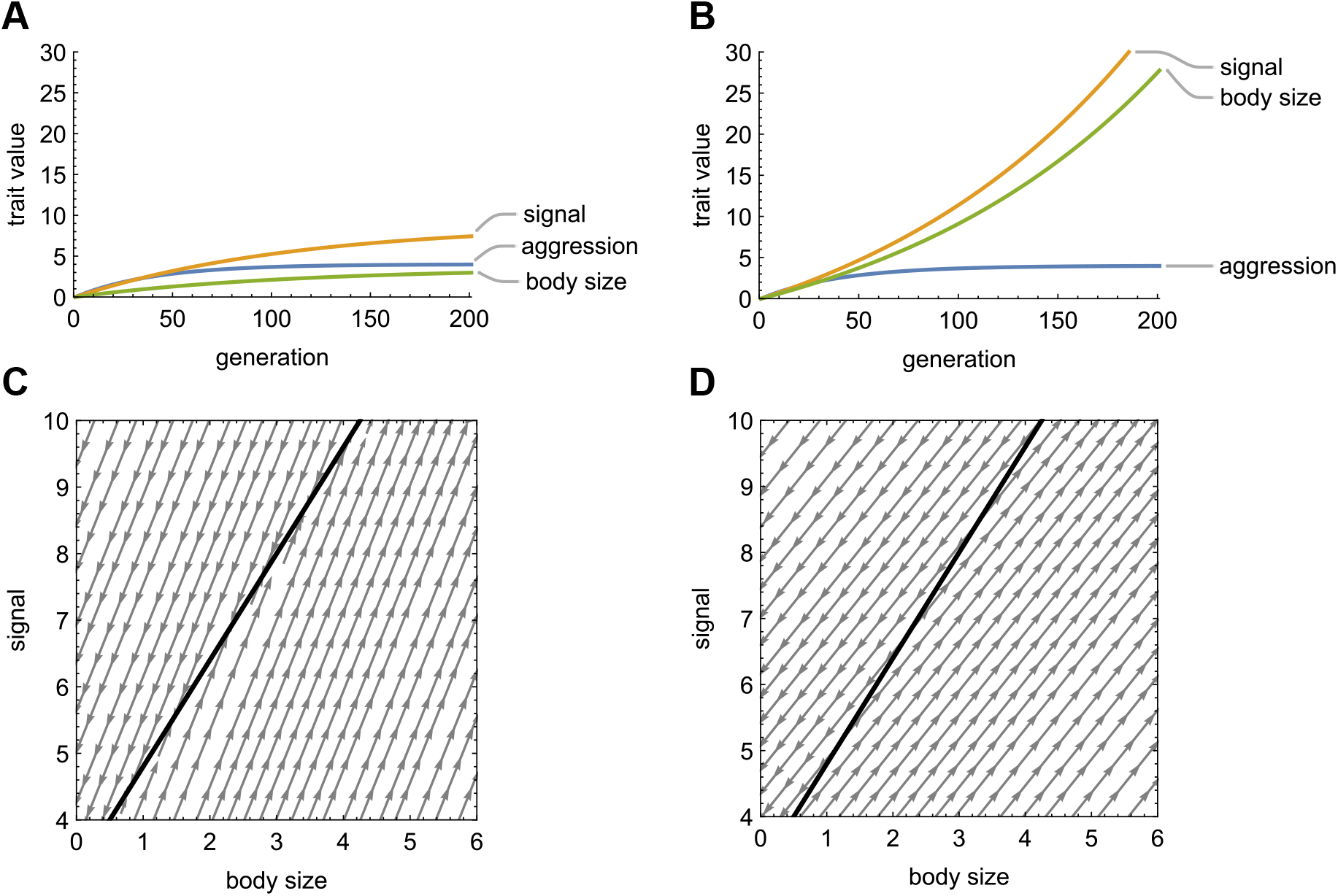
Evolution of a male signal, body size, and aggression in response to male-male competition. Panels A and B show evolutionary trajectories for each trait over 200 generations, and panels C and D show predicted lines of equilibria (heavy line) and their stability (gray arrows). In all panels, all three traits have the same genetic variance (*G* = 1), benefit (*b*_*A*_ = 0.2) and cost of aggression (*c*_*A*_ = 0.05), fitness cost deriving from the threat of a male’s opponent (*c*_*B*_ = 0.2), cost of signal size (*c*_*s*_ = 0.05; with naturally selected optimum *θ*_*S*_ = 0), and a responsiveness of aggression to body size (*δ*_*AS*_ = 0.4). The line of equilibria is calculated from Eq. 9a using these values. In panels A and C, signal size is weakly correlated with body size (*G*_*SB*_ = 0.4), while in panels B and D, the two traits are more strongly correlated (*G*_*SB*_ = 0.8). When the genetic correlation between signal size and body size is weak, all three traits reach equilibria (A), with equilibrium aggression predicted solely by costs and benefits. Signal size and body size reach a point on the predicted line of equilibrium (C) that differs depending on their starting values. When the genetic correlation is strong, aggression still reaches an equilibrium, but signal size and body size run away together (B), overshooting the predicted line of equilibria (D). As in Fisherian selection from female mate choice (Lande 1981), male-male competition can drive traits to runaway elaboration or extinction when the line of equilibria is unstable (D).

**FIGURE 2.**
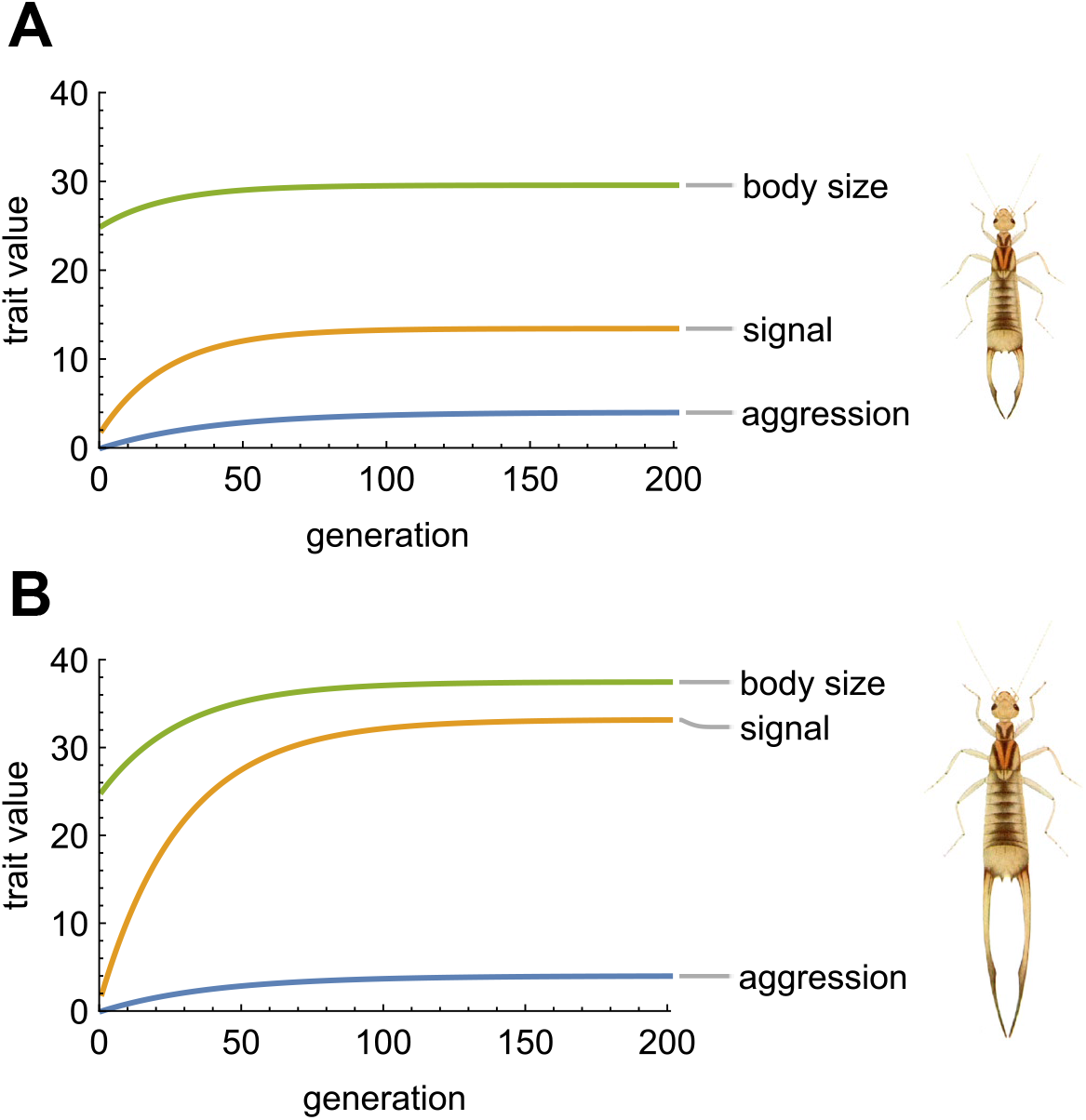
Stronger dependence of male aggressive behavior leads to more elaborate traits at equilibrium. Panel A illustrates a relatively weak influence of opponent signal on male aggression (*δ*_*AS*_ = 0.4), while panel B illustrates a stronger influence (*δ*_*AS*_ = 0.8). In each panel, we use starting values for traits relevant to the highly sexually dimorphic earwig *Labidura riparia*, which uses its forceps as a signaling trait and is shown to the right of each panel (drawing modified from Lucas 1920). Other parameters are the same as Fig. 1A. When the influence of opponent signal is weak (A), both body size and signal show a moderate evolutionary increase. When the influence is stronger (B), both body size and signal increase more, but the final signal size is much larger relative to body size. The highly elaborate elongate forceps in panel B may be found in other earwig species like *Forcipula gariazzi*.

Whether an evolving population will reach a predicted equilibrium (no runaway) or overshoot it (runaway) also depends on the rate of evolution of body size versus natural selection on signal size. Specifically, from Eq. 8a, for runaway evolution of signal size, body size must evolve fast enough so that 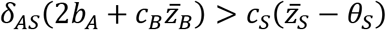). Because *b*_*A*_ and *θ*_*S*_ are constants, this occurs when 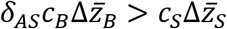, or equivalently:

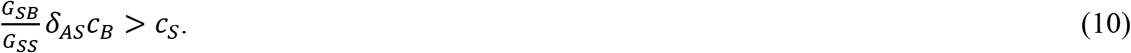

This result is also achievable by solving for the condition generating a negative eigenvalue of the Jacobian of 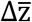, which indicates an unstable equilibrium (Lande 1981; Bailey & Kölliker 2019).

The first term in Eq. 10 is the regression of body size on signal size, which is typically large for weapons and signals (McCullough *et al.* 2016; Eberhard *et al.* 2018). As a regression coefficient, this term measures the degree to which body size can be predicted from signal size, which captures the logic of why the term measures signal honesty. In addition, Eq.10 indicates that runaway evolution of a signal is most likely to occur when three conditions exist: the signal is honest (*G*_*SB*_ is large and positive), it modifies aggressive behavior of social partners (*δ*_*AS*_ > 0), and aggression imposes a fitness cost that increases when opponents are larger (*C*_*B*_). Fig. 1 illustrates a scenario in which the predicted outcome (equilibrium or runaway) depends upon the value of the genetic covariance *G*_*SB*_.

## DISCUSSION

Our model provides explicit conditions for sexual selection arising from male-male competition to result in elaborate signals and runaway evolution. We model the origin of costs and benefits associated with male traits mediating male-male interactions using considerations from evolutionary game theory, which allows us to derive expressions for natural and social selection gradients that reflect the mechanistic properties of male contests (Eqs. 5-6). We incorporate these expressions for selection into a model of trait genetics based on the interacting phenotypes framework, which accounts for the influence of indirect genetic effects arising from interactions with an opponent (Eq. 2). Elaboration of a signal occurs whenever males adjust their level of aggression based on the signal of an opponent; i.e., *δ*_*AS*_ > 0 (Eq. 8a). This elaboration becomes runaway evolution when the signal is honest and when the cost imposed by aggression in an opponent increases with their body size (Eqs. 9a, 10; Figs. 1-2). In contrast, aggression always reaches an equilibrium, both because the fitness benefit of aggression is relative to that of the opponent and because of the fitness costs of escalated contests (Eq. 9c). Limits to runaway evolution of the signaling trait depend on the strength of natural selection opposing signal elaboration, which may arise through costs of producing or bearing the signal.

Our model does not specify the nature of the costs and benefits associated with aggression, the signaling trait, and body size (condition). These are important variables, likely ecologically contingent, and empirical work that quantifies these costs and benefits will provide context for the generality of our model. However, one of the strengths of this quantitative genetic modeling approach is that it provides predictions that are testable in natural populations. Specifically, we expect the evolution of elaborate signaling traits that resolve duels between males to evolve when three conditions are present. First, signals should be reliable predictors of body size or some other proxy of fighting ability. Indeed, such signal honesty, which is often characterized as positive allometry (McCullough *et al.* 2016; O’Brien *et al.* 2018) or a positive genetic correlation between size and signal (Clark & Moore 1995; McGlothlin *et al.* 2005; Laidre & Johnstone 2013), is a common feature of traits involved in male-male competition. Second, males must modify their behavior in response to their opponent’s signal. We assume that males increase their aggression when encountering an opponent with a smaller signal than their own and reduce their aggression when encountering an opponent with a larger signal. Such adjustment is common in species that resolve contests via limited fights or displays (Darwin 1871; West-Eberhard 1979, 1983; Emlen 2008, 2014). In our model, this phenomenon alters the relationship between genotype and phenotype, causing a net force of social selection to contribute to signal evolution (Eqs. 7a, 8a). Finally, we expect social selection to be imposed via the aggression of opponents. This selection becomes stronger as male body size or fighting ability evolves due to the threat of escalation of fights with large opponents. Mean level of aggression need not change if the threat escalates. Our model makes specific predictions for the signs of these gradients when selection on signal size, body size, and aggression can all be measured (Eqs. 5-6). Most crucially, our model predicts negative social selection gradients for both body size and aggression, which reflect the costs of escalated contests. In populations that are experiencing an evolutionary runaway, these gradients should become stronger as body size and signal size coevolve. Although few studies have measured social selection gradients, the limited evidence that exists supports the existence of negative social selection gradients imposed by competitors (Formica *et al.* 2011; Fisher & Pruitt 2019).

### Parallels to Lande’s model of female choice

The results of our model are conceptually analogous to Lande’s (1981) model of runaway sexual selection via female choice, suggesting some key parallels between the processes. Both our model and Lande’s, which was the first formal model of Fisher’s runaway process, result in lines of equilibria that may be stable or unstable depending on the genetic parameters. For the scenario of relative mate preference in Lande’s model, the line of equilibria for a male trait 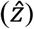 and a female preference 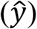 can be written:

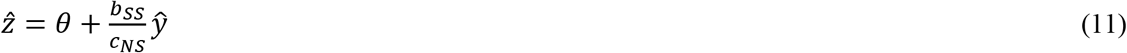

where *θ* is the naturally selected optimum, *b*_*SS*_ is the strength of sexual selection, and *C*_*NS*_ is the strength of natural selection. Eq. 11 directly parallels Eq. 9a from our model and emphasizes that in male-male competition, the force of sexual selection is provided not by direct female choice but by male body size (or some other measure of willingness or ability to engage in aggression). In male-male competition, the threat of the opponent (*C*_*B*_) leads to social selection, which is indirectly translated into evolutionary change in male signals via the parameter *δ*_*AS*_, measuring the dependence of aggression on relative signal size of two competing males.

Similarly, the condition for runaway evolution of male traits and female preference driven by mate choice in Lande’s model can be written:

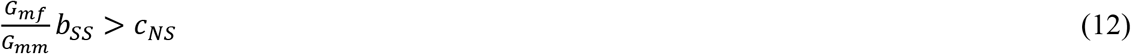

where *G*_*mf*_ represents the genetic covariance between male trait and female preference and *G*_*mm*_ represents genetic variance of the male trait. The condition in Eq. 12 directly parallels the condition in Eq. 10, emphasizing again that in male-male competition, *δ*_*AS*_*C*_*B*_ provides the force of social selection that indirectly leads to an evolutionary increase in male signal size. Both types of runaway evolution are driven by genetic covariance. In mate choice, runaway is driven by the covariance between the sexes that arises from choosy females mating with attractive males, but in male-male competition, this effect arises directly from signal honesty, i.e., the genetic covariance between a signaling trait and the threat (willingness or ability to fight) it signals. Moreover, if the mean level of aggression does not change, as when the aggression plateau is reached (Figs. 1-2), increasing costs during male-male competition are associated only with the increasingly elaborated signal. This may occur when limited fights settle contests (Maynard Smith & Harper 1988; Maynard Smith & Harper 2003). These are common conditions (Parker 1974; West-Eberhard 1983, 1984; Maynard Smith & Harper 1988; Andersson 1994), suggesting that runaway from male-male competition may occur frequently (McCullough *et al.* 2016; Rico-Guevara & Hurme 2019). Finally, the genetic covariance in Lande’s model arises from linkage disequilibrium that accumulates via nonrandom mating whereas ours reflects pleiotropy between body size and signal size. Thus, the genetic covariance driving runaway from male-male competition is likely to be much larger both because recombination efficiently erodes linkage disequilibrium and positive allometry (e.g., signals and body size) is common and reflects pleiotropy. Indeed, in models of mate choice where there is a pleiotropic relationship between direct benefits or good genes, runaway is difficult. In our model, however, the pleotropic nature of the honest signal leads to runaway.

There are many other models of mate choice in the literature, and a full comparison to them all is beyond the scope of this paper. Our goal here is simply to highlight potentially common features of runaway evolution, the most important is that runaway sexual selection by both male-male competition and female mate choice appears to be an evolving source of selection provided by the social environment. A more expansive comparison may well stimulate modifications or additions to the model we present here. In addition, there would be much to gain by combining studies of female mate choice and male-male competition to simultaneously test models of sexual selection (Hunt *et al.* 2009). This may be especially enlightening when traits serve as both ornaments and armaments (Berglund *et al.* 1996) or when mate choice opposes male-male competition (Moore & Moore 1999).

## Conclusion

Ritualized displays and elaborated signals associated with the potential for aggression are readily observed in nature and their importance often obvious and spectacular (Darwin 1871; Parker 1974; Maynard Smith & Harper 1988; Maynard Smith & Harper 2003; Emlen 2008, 2014). Yet the details of how these might evolve have been unclear. Previous game theory models have shown that overt aggression can be ameliorated by conventional signals (Parker 1974; Maynard Smith 1982; Maynard Smith & Harper 1988; Maynard Smith & Harper 2003; Rutte *et al.* 2006), and verbal models have proposed that signaling traits associated with male-male competition evolve exaggerated expression because social selection is intense (West-Eberhard 1979, 1983, 1984). Male-male competition may well result in intense selection (Maynard Smith & Brown 1986), as mating can be highly skewed toward one or a few males in a population (Darwin 1871; Andersson 1994; Shuster & Wade 2003), but this alone is insufficient to result in exaggerated traits. Our model shows that feedback between the behavioral and morphological traits mediating male-male competition create runaway evolution.

Sexual selection arising from male-male competition is prevalent and so the consequences of such selection important for understanding the generation of biological diversity. We hope our model stimulates empiricists in much the same way that the model of mate choice stimulated research on mating preferences. Ultimately, our understanding of the consequences of sexual selection arising from male-male competition will come from empirical research. Our hope is that this model helps direct and focus some of that research.

## SUPPORTING APPENDIX

## Fitness model for male-male interactions

To model the fitness consequences of an interaction between two males, we use a modification of the classic hawk-dove game, which involves competition over a resource (Maynard Smith & Price 1973, Maynard Smith 1982). In a traditional hawk-dove game, there are two strategies. Hawks tend to start fights, while doves tend to flee. When two hawks meet, a fight determines the outcome of the contest, with one hawk winning and receiving a resource of value *v*, while the other hawk pays a cost *C*, which may be related to injury or other costs of aggression. The average payoff for a hawk in a hawk-hawk encounter is thus 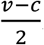. When two doves meet, they either divide the resource evenly or decide the contest without aggression and its associated costs, leading to an average payoff for each dove of 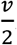. When a hawk meets a dove, the hawk wins the contest, receiving a fitness payoff of *v*, while the dove receives a payoff of 0, reflecting the fact that the dove neither wins the resource nor pays a cost for being aggressive. If *z* = 1 represents playing hawk and *z* = 0 represents playing dove, this fitness model can be written in terms of the phenotypes of two interactants expressing traits *z* and *z*′:

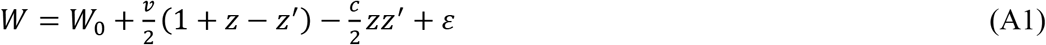

Following McGlothlin et al. (2022), we generalize this equation to a continuous scale, allowing *z*_*A*_, representing aggression, to take on any non-negative value. This means that individuals that are more aggressive (i.e., that employ a more hawkish strategy) will tend to win interactions and that the fitness cost of being aggressive will increase as both individuals become more aggressive (i.e., more hawkish). The fitness model in Eq. 4 differs from Eq. A1 by being expressed in terms of relative fitness (*w*) rather than absolute fitness (*W*). Thus, the translations between the parameters in the general hawk-dove fitness model and the fitness effects of aggression in Eq. (4) are

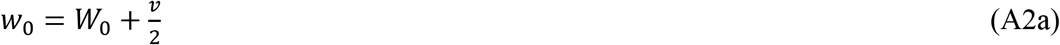

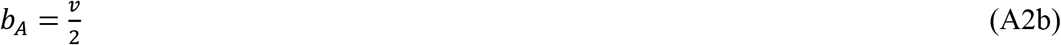

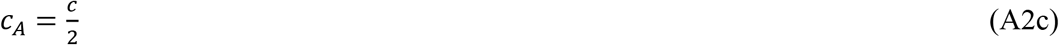

In words, *b*_*A*_ represents access to resources (here, mates) achieved by winning a contest, while *c*_*A*_ represents the costs paid by the loser of a contest when both individuals are aggressive. We also include two additional unique costs. We assume that the cost of an encounter is proportional to the body size (or fighting ability) of an opponent, resulting in a cost proportional to *c*_*B*_. Importantly, this cost is paid regardless of an individual’s own behavior. Second, bearing a signal larger than the naturally selected optimum imposes a cost proportional to *c*_*S*_, which is presumably paid outside the context of male-male encounters. Nonsocial and social selection gradients may be calculated from fitness functions such as Eq. A1 and Eq. 4 by taking the partial derivatives of relative fitness with respect to focal and social phenotypes, respectively (McGlothlin *et al.* 2022). These derivatives are then evaluated at the population mean. For linear fitness functions, selection gradients will be constant, but quadratic and higher-order functions may lead to selection gradients that change with the population mean.

## General equation for response to selection

Here, we develop a general model for evolution when trait expression depends upon the difference between a focal individual’s own traits and traits of another individual encountered in the context of a social interaction. This model is directly applied to male-male contests in the main text and may be useful for considering many other types of social interactions. First, consider a vector of traits (**z**) whose expression can be decomposed into three components: a vector of additive genetic effects (**a**), a vector of environmental effects (**e**), and a social response term that depends on the difference between the traits of the focal individual and those of a social interactant (**z**′):

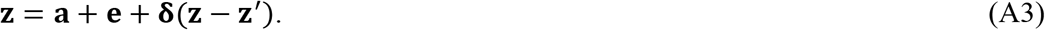

The matrix **δ** consists of components *δ*_*ij*_ that translate the effect of differences in trait *j* into expression of trait *i*. Similarly, the phenotype of the social partner can be written as

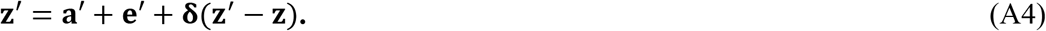

As we show below, because the term **δ**(**z** − **z**′) in Eqs. A3-A4 contains phenotypes of both individuals, it consists of a combination of direct and indirect genetic effects.

To calculate a response to selection for traits expressed as in Eq. A3, we first solve for the multivariate phenotypic mean. Assuming that environmental effects have a mean of zero, the trait mean is

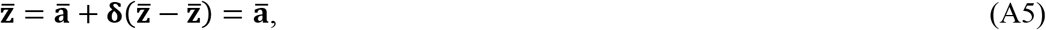

which means that the population trait mean will depend only on the mean additive genetic value. The vector of total breeding values (**A**), which represents the genetic contribution to the population mean and is used to calculate evolutionary responses to selection, is equivalent to the vector of additive genetic effects (**a**). Next, to derive an explicit definition of the phenotype, we first use substitution to write

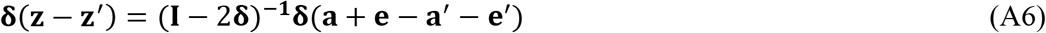

where **I** is the identity matrix. After some algebra, Eq. A6 allows us to write explicit definitions of the two phenotypes as

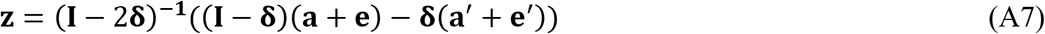

and

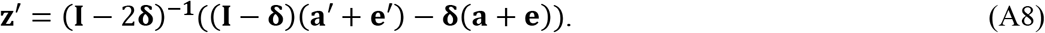

Response to selection can then be calculated following (McGlothlin *et al.* 2010) as

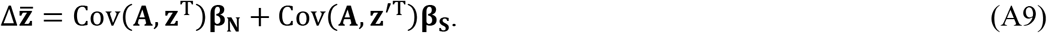

Substituting Eqs. A7-A8 into Eq. A9 and simplifying yields

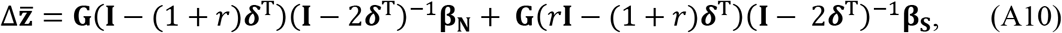

where **G** is the additive genetic (co)variance matrix and *r* is relatedness. Eq. A10 can be used to generate the specific model in the text by setting parameters as

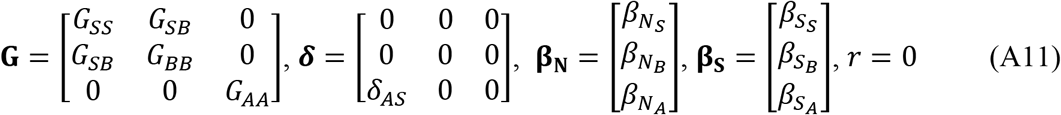

and multiplying the result by 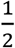 to indicate selection acting only on males.

## Relationship to indirect genetic effects models

The model described above differs from previous models of indirect genetic effects in that phenotypes may be adjusted in relation to both the phenotypes of a social partner and other phenotypes of the focal individual. The standard model of phenotypic expression used in trait-based genetic effects models (Moore *et al.* 1997a) is:

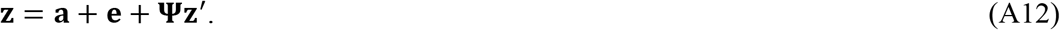

The relationship between the two models can be seen by adding an additional term to the standard model:

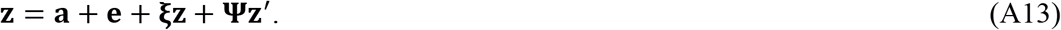

The term **ξz**, which is similar to developmental interaction effects (Wolf *et al.* 2001), contains a conditional modification of the direct genetic effect in response to other phenotypic traits of the same individual, while the term **Ψz**′ contains indirect genetic effects (Moore *et al.* 1997a). When **ξ** = **0**, Eq. 13 is equivalent to Eq. A12, and when **ξ** = **−Ψ** = **δ**, Eq. A13 is equivalent to Eq. A3.

Incorporating conditional direct genetic effects is a way to mechanistically represent genetic covariances in a quantitative genetic model (Wolf *et al.* 2001). Combining such effects with indirect genetic effects allows exploration of a wide variety of models of phenotypic adjustment, including behavioral modification, in evolutionary quantitative genetic models. For full generality, we give the equation for multivariate response to selection derived from Eq. A13. First, the vector of total breeding values derived from the trait mean is

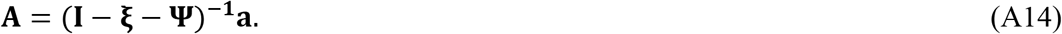

By substituting Eqs. A13-A14 into Eq. A9,

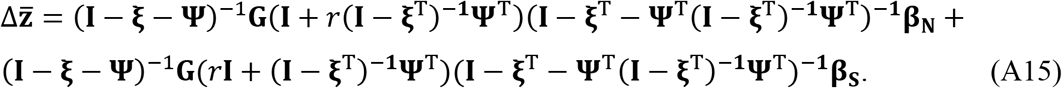

This equation is unwieldy in its multivariate form, but one can follow the approach we take here for male-male competition and use Eq. A15 to generate much simpler models of the evolution of systems of responsive traits given assumptions about fitness and trait expression.

## Extension to multiple opponents

Our results can be extended to incorporate interactions with multiple opponents. Suppose that a male encounters opponents sequentially. The phenotype of a focal individual averaged over all encounters:

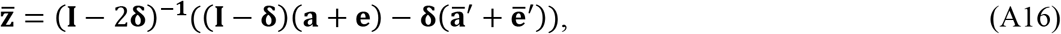

and the average of his social partners:

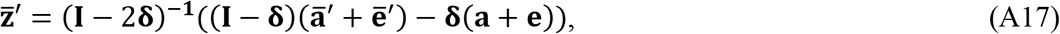

where an overbar is now taken to indicate an average over an individual male’s encounters. If we assume that the fitness effects of individual encounters accrue additively, then Eq. A10 may still be used to predict the evolution of the population mean, with **β**_**N**_ and **β**_**S**_ now representing vectors of partial regression slopes of fitness on focal individual mean and social group mean phenotypes, respectively. Specific fitness models may be substituted into Eq. A10 in the same way as for models of single pairwise interactions.

## ACKNOWLEDGMENTS

We thank our collaborators, students, and postdoctoral scientists who over the years provided discussions and comments on various earlier versions of this idea and model. This work was supported by grants from NERC (UK, NE/C510659/1 to JBW) and NSF (USA, IOS-1354358 to AJM, DEB-1457463 to JWM).

